# Synaptic inhibition in the accessory olfactory bulb regulates pheromone location learning and memory

**DOI:** 10.1101/2024.09.08.611942

**Authors:** Shruti D. Marathe, Sanyukta Pandey, Devesh Rawat, Susobhan Das, Nixon M. Abraham

## Abstract

Pheromone signaling is pivotal in driving the social and reproductive behaviors of rodents. Learning and memorizing the pheromone locations involve olfactory subsystems. To study the neural basis of this behavior, we trained female heterozygous knockouts of GluA2 (AMPAR subunit) and NR1 (NMDAR subunit), targeting GAD65 interneuron population, in a pheromone location learning assay. We observed impaired memory of pheromone locations on early and late recall periods, pointing towards the possible role of ionotropic glutamate receptors (iGluRs), and thereby the synaptic inhibition in pheromone location learning. Correlated changes were observed in the expression levels of activity-regulated cytoskeletal (*Arc)* protein, which is critical for memory consolidation, in the associated brain areas. Further, to probe the involvement of the main and accessory olfactory bulbs (MOB and AOB) in pheromone location learning, we knocked out GluA2 and NR1 from MOB and/or AOB neuronal circuits by stereotaxic injection of Cre-dependent AAV5 viral particles. Perturbing the inhibitory circuits of MOB and AOB or AOB-alone resulted in the pheromone location memory deficits. These results confirm the role of iGluRs and the synaptic inhibition exerted by the interneuron network of AOB in regulating learning and memory of pheromone locations.

## Introduction

The vomeronasal system, one of the subsystems of rodent olfactory system, plays a critical role in pheromone information processing (1, 2). The neuronal circuitry, starting at the vomeronasal organ (VNO) and passing through the accessory olfactory bulb, vomeronasal amygdala, and the preoptic area of the hypothalamus, is involved in this process, which is vital for the survival and mating behavior of animals (3–7). While terrestrial rodents encounter pheromones streaked on objects of different shapes and sizes, their whisker system is involved in sampling the features of objects. This multimodal association in pheromone location learning has been proven previously (8). Here, we addressed the neural mechanisms of pheromone location learning and memory by focusing on the main and accessory olfactory bulb (MOB and AOB) neuronal circuits.

Male mice mark their territories through urine depositions in many social contexts (9). It has been found that non-volatile components of mouse urinary pheromones are processed by the MOB circuits and the volatiles by the AOB circuits. While the VNO type 1, type 2, and formyl peptide receptors sense the non-volatile components, trace amine-associated receptors expressed on the main olfactory epithelium are activated by the volatile pheromones (3, 10–12). This information is further received and processed at the AOB and MOB by the projection neurons (13, 14). The inhibitory network of MOB and AOB, which forms dendrodendritic synapses with the projection neurons refines the odorant/pheromone information before being conveyed to the higher centers (15–18).

The vomeronasal sensory neurons (VSNs) expressing the same vomeronasal receptors project to multiple glomeruli in the AOB. This organization is different from that of MOB (19–21). Unlike in MOB, the projection neuron cell bodies are dispersed in AOB, and their primary dendrites target multiple glomeruli (22, 23). AOB projection neurons have been shown to be selective for the sex of urine donors, and the inhibitory network is critical in shaping this selectivity (24, 25). The inhibitory network of the AOB is constituted by GABAergic periglomerular interneurons and granule cells (GCs) (26, 27). There are several morphologically and physiologically distinct classes of interneurons in the AOB. Among these, GAD65-expressing ones represent a significant population (17). The projection neurons release glutamate, which activates ionotropic glutamate receptors (iGluRs) on GCs, and in response, they release GABA, which causes inhibitory feedback on the projection neurons (28).

To probe the neural mechanisms of pheromone location learning, we modulated the activity of the MOB and AOB inhibitory networks by targeting iGluRs expressed on GAD65-expressing inhibitory interneurons (29). After the conditional deletion of the GluA2 subunit of AMPARs and the NR1 subunit of NMDARs, we trained female heterozygous knockouts (KOs) of GluA2 and NR1 in a pheromone location learning assay. We observed impaired memory of pheromone locations in both early and late recall periods, indicating the possible role of iGluRs and, thereby, synaptic inhibition in pheromone location learning and memory. Correlated changes were observed in the expression levels of activity-regulated cytoskeletal (*Arc)* protein, which is critical for memory consolidation, in the associated brain areas. To study the involvement of MOB and AOB microcircuitry in learning pheromone locations, we specifically knocked out NR1 and GluA2 from MOB and/or AOB neuronal circuits by stereotaxically injecting Cre-dependent AAV5 viral particles. Perturbing the neuronal circuitry of MOB and AOB or AOB-alone resulted in the loss of pheromone location memory. These results confirm the role of iGluRs and the synaptic inhibition exerted by the interneuron network of AOB in controlling the learning and memory of pheromone locations.

## Materials and Methods

### Animals

A total of 37 control C57BL/6J female mice, aged 8-12 weeks, were utilized for these experiments (experiment-wise breakup of animals is mentioned in the corresponding data). In addition to wild-type animals, transgenic mice were obtained by crossing the following genotypes obtained from Jackson Laboratory unless otherwise mentioned.

GluA2^2Lox^: GluA2^2Lox^ (Gria2) was obtained from the Heidelberg University, Germany

GluA2^GAD65(+/-)^ knockout: GluA2^2Lox^ with B6N.Cg-Gad2^tm2(cre)Zjh^/J for the knockout of GluA2 in GAD65+ve neurons. NR1^2LOX^: NR1^2Lox^ (GRIN1) was obtained from the Heidelberg University, Germany

NR1^GAD65(+/-)^ knockout: NR1^2Lox^ with B6N.Cg-Gad2^tm2(cre)Zjh^/J for the knockout of NR1 in GAD65+ve neurons. GAD65-GCaMP6f: B6N.Cg-Gad2^tm2(cre)Zjh^/J with B6;129S-Gt(ROSA)26Sor^tm95.1(CAGGCaMP6f)Hze^/J for the expression of GCaMP6f in GAD65+ve cells.

A total of 40 NR1^GAD65(+/-)^ and 28 GluA2^GAD65(+/-)^ female mice aged 8–12 weeks were used in this study. 18 male mice (non-littermates GluA2Lox, NR1Lox) aged 10–16 weeks were used to collect urine and soiled bedding. The control groups included GluA2Lox/NR1Lox.

### Ethical Approval

The experimental procedures used in this study are approved by the Institutional Animal Ethics Committee (IAEC) at IISER Pune and the Committee for the Control and Supervision of Experiments on Animals (CCSEA), Government of India (animal facility CCSEA registration number 1496/GO/ReBi/S/11/CCSEA). The usage of animals in this study was approved under protocol number IISER_Pune/IAEC/2016_01/001.

### Calcium Imaging

Calcium imaging was performed, as explained previously (18, 29). In brief, for calcium imaging, 5 naïve adult females, 8 to 12 weeks of age, were used. The animals were anesthetized with a mixture of ketamine and xylazine, and a 1mm diameter cranial window was created at the posterior end of the right hemisphere of the OB using a dental drill. As a precaution, a blunt Hamilton needle was lowered 1.2 mm ventral to the dura in the AOB and kept for 5 minutes to create a path before insertion of the GRIN lens. The 1.2 mm protruding GRIN lens was lowered vertically until it reached the AOB. To stabilize the lens assembly to the skull surface, a mixture of cyanoacrylate gum and acrylic dental cement was used. A headpost was implanted behind the implanted lens, and animals were given a month to recover. Ca2+ imaging was performed after the recovery period in awake head-restrained conditions. The activity of GAD65 interneurons was recorded in response to 0.4LPM airflow and male urine for 40 trials each before starting the training. Animals were then subjected to 4 days of testing (ITD) and 15 days of training (TrD) and calcium activity was recorded again for 0.4LPM airflow and male urine in two consecutive sessions (40 trials/animal/stimulus condition) following last training day. Mice were perfused using Dulbecco’s PBS, following the imaging sessions post-training, brains were dissected and kept overnight for fixation in 4%PFA. Coronal OB sections of 80 µm were taken and scanned under the confocal microscope to confirm the lens path. Further, the data was analyzed using custom-written Python scripts. A heatmap and selected individual traces were plotted along with average plots for ΔF/F.

### Genotyping

NR1Lox and GluA2Lox mice were crossed with GAD65-Cre mice to produce NR1^GAD65(+/-)^ and GluA2^GAD65(+/-)^ heterozygous knockout (KO) animals respectively. Tail samples of the F1 generation mice were collected and their DNA was isolated using a KAPA DNA extraction kit according to manufacturer’s instructions. DNA quality and concentration were measured using NanoDrop (Thermo Fisher Scientific). The genotype was confirmed by Polymerase Chain Reaction (PCR), followed by Gel Electrophoresis. The sequence of primers used, and the reaction conditions are shown in Table 1.

### Protein quantification

To quantify the drop in the protein levels of NR1 and GluA2, western blotting was performed. The following groups of animals were used for western blotting: control (NR1^2Lox^, GluA2^2Lox^), NR1^GAD65(+/-)^, and GluA2^GAD65(+/-)^ animals [n (number of animals) = 3 each, N (number of replicates) = 2]. Animals aged 6 to 10 weeks were utilized. The animals were anesthetized in a CO_2_ chamber and then decapitated. Both OBs were dissected and flash-frozen immediately. The samples were stored at-80^0^C until further use. OB lysates were prepared in RIPA buffer (50 mM Tris HCl, pH 7.4, 150 mM NaCl, 1% Triton X-100, 0.5% Sodium deoxycholate, 0.1% SDS, 1 mM EDTA, 10 mM NaF), supplemented with a complete protease inhibitor (Roche Cat # 04693116001). The BCA assay was performed to quantify the total protein concentration using a Pierce BCA protein assay kit (Thermofisher Cat # 23225). Approximately 20 μg of protein from each sample was loaded onto a 12% SDS-Polyacrylamide Gel (SDS-PAGE). After separation, the proteins were electrophoretically transferred to an Immobilon-P PVDF membranes (Millipore Cat # IPVH00010) through methanol buffer transfer. To selectively block non-specific sites, the membrane was incubated in 5% non-fat dry milk/Tris Buffer Saline (TBS)-Tween 20 for 1 hour at room temperature. The membranes were probed with primary antibodies: anti-NR1 antibody (Millipore Cat # AB9864R), anti-GluA2 antibody (Millipore Cat # AB1786-l), and anti-GAPDH (Sigma Cat # G9545) at 1:1000, 1:250 and 1:5000 dilutions, respectively, for approximately 20 hours at 4^0^C. The blots were washed with 0.1% TBST three times for 15 minutes each. The membrane was subsequently incubated in a 1:5000 dilution of peroxidase-conjugated AffiniPure Goat anti-rabbit IgG (Jackson ImmunoResearch Cat # 111-035-003) for 1 hour at room temperature. Finally, the protein bands were detected using Clarity ECL Western Blotting Substrate (BioRad Cat # 1705061), and the images were digitally acquired using the ImageQuant LAS 4000.

### Multimodal Pheromonal location learning apparatus

A non-reflective, black-painted plexiglass apparatus measuring 60 cm (L) x 30 cm (W) x 15 cm (H) was utilized for multimodal pheromone location learning. The setup was divided into three equal zones, which could be isolated using removable plates. At the opposite ends of the apparatus, there were chambers measuring 10 cm x 10 cm x 15 cm. These chambers were centrally located and had removable plates with equidistant holes, either 10 mm or 5 mm in diameter. In one of the chambers, called the urine chamber, 100 μl of male urine (opposite-sex pheromone) was placed in a 55 mm Petri dish guarded by a plate with 10 mm holes, while 100 μl of water (a neutral stimulus) was placed in the other chamber, called the water chamber, guarded with plate having 5 mm holes. The plates with different hole-sizes were counterbalanced for different animals. The odor and pheromone released through the holes in the plate, restricting the access of female mice to the odors/pheromones on only one side of the chamber. Two different identical setups were used for the training and memory sessions (8).

### Multimodal Pheromone Location Learning Paradigm

The pheromone location learning and memory assay was carried out, as explained previously (8). The assay consists of three phases: 1) the Initial Testing Days (ITDs), 2) the Training Days (TrDs), and 3) Memory Day (MD). The ITD was performed for an initial four days. During ITDs, the female mouse was allowed to roam in the setup unrestricted for 10 mins, and the activity was video recorded. The ITD is done to check if any female has an innate preference towards the male urine chamber over the water chamber. The setup was rotated by 180^0^ daily to avoid any bias towards spatial location throughout the ITDs and TrDs. During ITD, male urine (MU) and water (NS) are kept in the respective chambers. Soiled bedding (SB) from the male cage from which urine has collected is also spread at the front of the male urine chamber. This ensures volatiles as well as non-volatiles are emanating from the male pheromones.

During each experiment, female mice were counterbalanced towards a particular hole size on the plate (e.g., half female mice had a 10 mm hole-sized plate in the urine chamber and a 5 mm hole-sized plate in the water chamber. For the rest of the females, the urine chamber had a 5 mm hole-sized plate, and the water chamber had a 10 mm hole-sized plate). This counterbalancing was done to avoid any predisposition of female mice towards a particular hole size.

Following ITD 4, TrDs were conducted for the next 15 days. During the TrDs, female mice were confined to both chambers for 15 minutes each, alternating after every 5 minutes in urine and water chambers. This was continued for 30 min every day for 15 days. Every day, the setup was rotated by 180^0^. During each training day, fresh MU, SB and water were used. Memory was tested on a different setup which was identical to the training setup. This was done to avoid animals sensing any remnant cues in the setup.

### Behavior quantification

Two parameters were quantified in pheromone place assay, 1) ‘Time spent’ near Zone 1 vs Zone 2, and 2) ‘number of active attempts’ done towards respective plates. EthoVision software (Noldus Information Technology, version 8.5 XT) was used to quantify ‘time spent’. The nose-point feature was used for quantification. The number of active attempts suggested nose pokes done by the animal. Active attempts were counted manually.

### Immunohistochemistry

After memory days 7 and 15, animals were sacrificed and transcardially perfused with 1X PBS and 4% paraformaldehyde within 30 min of the experiment to quantify the expression of Activity-regulated cytoskeletal (Arc) (Synaptic Systems, Catalogue # 156003) in the olfactory bulb (main and accessory olfactory bulb), hippocampus and somatosensory cortex (SSC). Brains were kept in 4% PFA overnight at 4^0^C. A brief wash with 1X PBS was given, and the brains were transferred to 30% sucrose which acted as a cryoprotectant. Once the brains were submerged in the sucrose solution, they were embedded in a block using freezing media. 50 μm thick coronal sections were cut using a cryotome. We selected every 6^th^ section of OB, hippocampus, and SSC for staining purposes. The sections were washed with 1% triton in 1X PBS for 10 min. Then the sections were incubated in a blocking buffer (5% BSA in 0.5% triton and 1X PBS) for 1.5 hours at room temperature. Primary antibody incubation (1:750 μl of anti-Rabbit Arc antibody in 0.5% BSA in 0.1% triton and 1X PBS) was done overnight at 4^0^C. Three washes with 1X PBS were done for 15 min each. During the last wash DAPI (4’,6’-diamidino-2-phenylindole) was added to visualize nuclei under the microscope. Secondary antibody incubation (anti-Rabbit Alexa-fluor 488 1:1000) was done for 2 hours at room temperature. Three washes with 1X PBS were done for 10 min each.

For c-Fos antibody staining, 50 μm thick sections were incubated with blocking buffer (5% NGS, 2.5% BSA in 0.5% triton in 1XPBS) for 2 hours at room temperature. The anti-c-Fos antibody (Rabbit anti-cFos antibody, 2250 S, Cell signaling technologies) was used at a dilution of 1:500 in blocking solution and incubated overnight at 4°C. A secondary antibody (Anti-rabbit Alexa fluor 594, Jacksons Immunoresearch,111-585-003) was used at 1: 1000 dilution for 1.5 hours at room temperature. The NeuN antibody staining was performed as described before (18). Briefly, the sections were washed with 0.1% triton in 1XPBS for 10 min. Then, the sections were incubated in a blocking buffer (7.5% Normal Goat Serum, 2.5% BSA in 1% triton, and 1X PBS) for 3 hours at room temperature. Primary antibody incubation (1:1000 ul of anti-Chicken NeuN antibody, ABN91, Merck in 7.5% NGS,2.5% BSA in 1% triton and 1 X PBS) was done overnight at 4^0^C. Three washes with 1 X PBS were done for 15 min each. A secondary antibody (Anti-chicken Alexa Fluor 647, 03-605-155, Jackson’s Immunoresearch, 1:1000) was used for 2 hours at room temperature. Three washes with 1 X PBS were done for 10 min each. The sections were mounted on slides and were light-protected with a Vectashield. The sections were imaged using a Leica SP8 confocal fluorescent upright microscope. Imaging was done under 20X.

### Confocal imaging and cell count quantification

c-Fos and Arc expressing cells across OB, AOB, hippocampus, and SSC were imaged using a Leica SP8 confocal microscope. 4 to 6 sections per brain area per mouse were used for quantification. The cell quantification was done using Imarisx64 and Huygens professional software (Bitplane, Oxford Instruments). For quantification, each section with a Z-stack of 1 μm step size was used. To reduce the background noise, A constant Gaussian filter was applied to every stack. The threshold used to count the c-Fos and Arc-expressing cells was automatically set by the software Imaris. False positive counts were removed manually. The total cells counted for each brain region was equal to the sum of the cells counted per stack. The total volume was obtained by multiplying the thickness of the stack with the XY diameter of the image. The final number of c-Fos and Arc positive cells counted was /1 mm^3^.

### Stereotaxic surgeries

Female mice aged 6 to 8 weeks were injected with Cre-viral particles (pAAV.CMV.HI.eGFP-Cre.WPRE.SV40, titer: 7 x 10^12^ vg/ml, Addgene), using a BENCHMARKTM stereotaxic instrument (myNeurolab, St. Louis, MO) and LEICA MZ6 stereo microscope (Leica Microsystems, Germany). MOB-specific injections were done as described previously (15). AOB-specific injection coordinates were decided based on the previous reports (30). In brief, the optimization of injection coordinates was done after a few fluorescent dye injections. The injection pipette tip was positioned at the bregma, and all axes were set to zero. The pipette tip was then moved to the anterior side near the intersection of the sagittal suture and infra-cerebral vein, where the Z coordinate was adjusted according to the Z coordinate at bregma. The coordinates were made zero across the three axes. Based on this local reference point, injection coordinates were decided. A total of two injections were done in each bulb. A resting period of 5 – 7 min was given after each injection to avoid backflow of the viral particles. At each spot, around 100 – 150 nl of solution was injected. The animals were allowed to recover for the next three weeks before starting with the behavioral experiment.

## Data and Statistical analysis

GraphPad Prism 9, Microsoft Excel, and Python were used for all statistical analyses in this study. The data is presented as Mean ± SEM. To determine p-values and test for statistical significance, we used the student’s t-test, one-way and two-way ANOVA, and associated post-hoc tests.

## Results

### Inhibitory networks of the main and accessory olfactory bulb are involved in pheromone location learning

Learning and memorizing the information about pheromone locations is critical for the reproductive success and survival of animals. To study the pheromone location learning abilities of mice, we developed a pheromone location learning and memory assay as described before (8). We trained a batch of wild-type female mice to associate specific hole sizes on the chamber plate with zones containing opposite-sex urine vs. neutral stimulus, water (Figure 1A). To investigate the role of olfactory bulb circuits in such learning, we compared the c-fos (a neuronal activity marker) expression pattern in a naive cohort of home-cage control mice and the trained group (Figure 1A). Colocalization of c-fos expression with GAD65-expressing neurons of MOB and AOB shows the involvement of inhibitory circuits in such learning (Figure 1 A2). We observed a significantly higher number of c-fos-expressing cells in the interneuron layers of OB, indicating the plausible involvement of the inhibitory network in pheromone location learning (Figure 1A3, Two-way ANOVA, followed by Tukey’s multiple comparisons test, p = 0.03).

**Figure 1:**
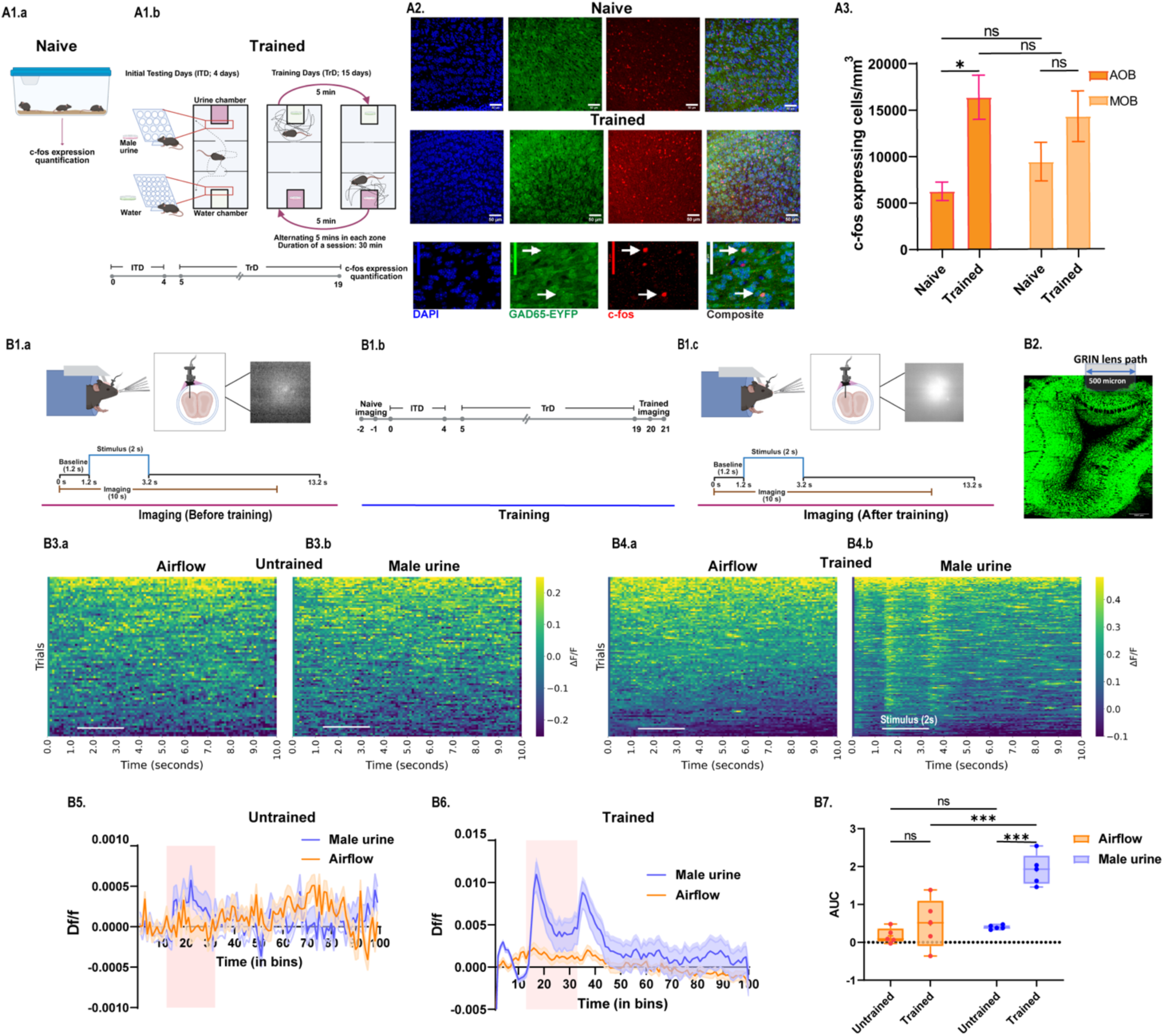
Inhibitory network of accessory olfactory bulb is involved in pheromone location learning. A1: A1a. Representation of two groups of GAD65-EYFP transgenic mice, i.e., naive (home cage control) and trained cohort used, respectively, for the c-Fos quantification and colocalization. A1b. Represents the pheromone location learning assay and timeline of training. A2: Representative confocal images showing DAPI (blue), c-Fos (red), and GAD65-EYFP (green) expressing cells in OB of naive (home-cage control) and trained groups of mice. A3. Bar graphs comparing c-Fos expression in AOB and MOB of naive and trained cohorts. We observed a significant increase in c-Fos expressing cells in AOB post-training day 15 compared to naive home cage controls. Two-way ANOVA p (Training status) = 0.005, followed by Tukey’s multiple comparisons test, AOB, Naive vs Trained p= 0.03, MOB, Naive vs Trained p = 0.09, N_Naive_, N_Trained_ = 3, respectively). B1a. Experimental setup to record Ca2+ dynamics under awake head-restrained conditions in untrained condition. Illustration representing micro-endoscopic calcium imaging and an example frame of Ca2+ response from the GAD65-expressing interneurons of AOB in response to male urine. Female mice were implanted with a metallic head-plate to facilitate restraining and a cylindrical GRIN lens to image Ca2+ dynamics. Scheme of stimulus presentation and imaging for the micro-endoscopic imaging from AOB is presented in the lower panel. B1b. Diagrammatic representation of the pheromonal location learning experimental timeline. Same group of mice was used for calcium imaging before (Untrained) and after training (Trained) on the pheromonal location learning assay. B1c. Experimental setup to record Ca2+ dynamics under awake head-restrained conditions after training. Illustration depicts micro-endoscopic calcium imaging and an example frame of Ca2+ response from the GAD65-expressing interneurons of AOB in response to male urine in trained mice. Scheme of stimulus presentation and calcium response recording is presented in the lower panel. B2. Coronal section showing GRIN lens path in mouse AOB (horizontal scale bar = 200 µm) B3.a, B3.b, B4.a & B4.b. Pseudo-colored heatmap showing Ca2+ responses in the GAD65-expressing interneurons of AOB in response to airflow (0.4LPM) and male urine exposure for 2 seconds (stimulus duration marked with white horizontal line) in untrained and trained conditions, respectively. B5, B6. Figure depicting changes in fluorescence, i.e., Df/f, in response to the airflow (0.4LPM) and male urine in untrained and trained conditions, respectively. B7. Bars comparing AUC values (area under the Df/f curve) during the stimulus window for airflow (0.4 LPM) and urine in untrained and trained conditions. Notably, trained condition exhibited a remarkable increase in the calcium activity for urine exposure compared to the airflow. Two-way ANOVA p (Training status X stimulus) = 0.0009, followed by Tukey’s multiple comparisons test, Airflow: Untrained vs Trained p = 0.08; Male Urine: Untrained vs Trained p = 0.0003; Untrained: Airflow vs Male Urine, p = 0.2; Trained: Airflow vs Male Urine, p = 0.0004, N = 5 mice). * p < 0.05, ** p < 0.01, *** p < 0.001, **** p < 0.0001, ns p > 0.05

To validate the involvement of GAD65-expressing interneurons in processing pheromonal cues and pheromonal location learning, we conducted calcium imaging experiments using transgenic female mice expressing GCaMP6f in GAD65-expressing interneurons, which comprise majority of the interneuron population in MOB and AOB, before and after training (Figure 1B). The GRIN lens was implanted in the AOB of GAD65-GCaMP6f transgenic female mice (Figure 1 B4). As a control, calcium activity was also recorded in response to airflow used to deliver pheromonal volatiles (0.4 liter per minute, LPM). This population activity was measured in awake, head-restrained condition. The fluorescence change (ΔF/F_0_) was quantified for 40 trials across 5 females in untrained and trained conditions (Figure 1 B3-B6). Calcium responses were observed from the interneuron population, indicating their involvement in processing the pheromonal information (Figure 1 B5-B6). Further, on quantifying the calcium responses post-training day 15, we observed a marked increase selectively for the male urine but not for airflow stimulation, proving the involvement of AOB inhibitory network in pheromone location learning and memory (Figure 1 B7, Two way ANOVA followed by Tukey’s multiple comparisons test, Untrained male urine vs. Trained male urine, p = 0.0003, Trained airflow vs. Trained male urine, p = 0.0004, n = 5).

### Ionotropic glutamate receptor functions of GAD65-expressing inhibitory interneurons are necessary for *pheromone location learning and memory*

To study the involvement of inhibitory interneurons, we used the transgenic mice in which ionotropic glutamate receptor functions were modified specifically in the GAD-65-expressing ones. To probe if the genetic perturbations result in any anxiety and despair-like behaviors, we carried out elevated plus maze, forced swim, and tail suspension tests. We did not observe any of these behavioral phenotypes in the genetically modified animals (Figure S1). Upon confirming no other behavioral disorders due to the genetic perturbations, we performed a pheromone detection assay using Controls, NR1^GAD65(+/-)^ and GluA2^GAD65(+/-)^ female mice. Animals were allowed to explore an arena where the soiled bedding (SB) with male urine was kept in a petri dish at the center of the arena. Time spent in the center zone of an empty chamber (open arena) similar to pheromone detection assay was taken as a control for comparison. Animals across groups (i.e. Controls, NR1^GAD65(+/-)^ and, GluA2^GAD65(+/-)^) spent significantly more time exploring the pheromone-containing center zone compared to the empty center-zone, suggesting intact detection abilities in the knock-outs (Figure 2A1-B5, Two-way ANOVA, followed by Tukey’s multiple comparison test, B5; Control Male urine - Control Open arena, p = 0.013, NR1^GAD65(+/-)^ Male urine - NR1^GAD65(+/-)^ Open arena, p = 0.001** and GluA2^GAD65(+/-)^ Male urine – GluA2^GAD65(+/-)^ Open arena, p = 0.009, N= 7-10). Additionally, time spent near the pheromone-containing chamber did not differ significantly across different groups Controls, NR1^GAD65(+/-)^ and GluA2^GAD65(+/-)^ female mice (Figure 2 B5: Two-way ANOVA, Tukey’s multiple comparison tests, p > 0.05, n = 7-9).

**Figure 2:**
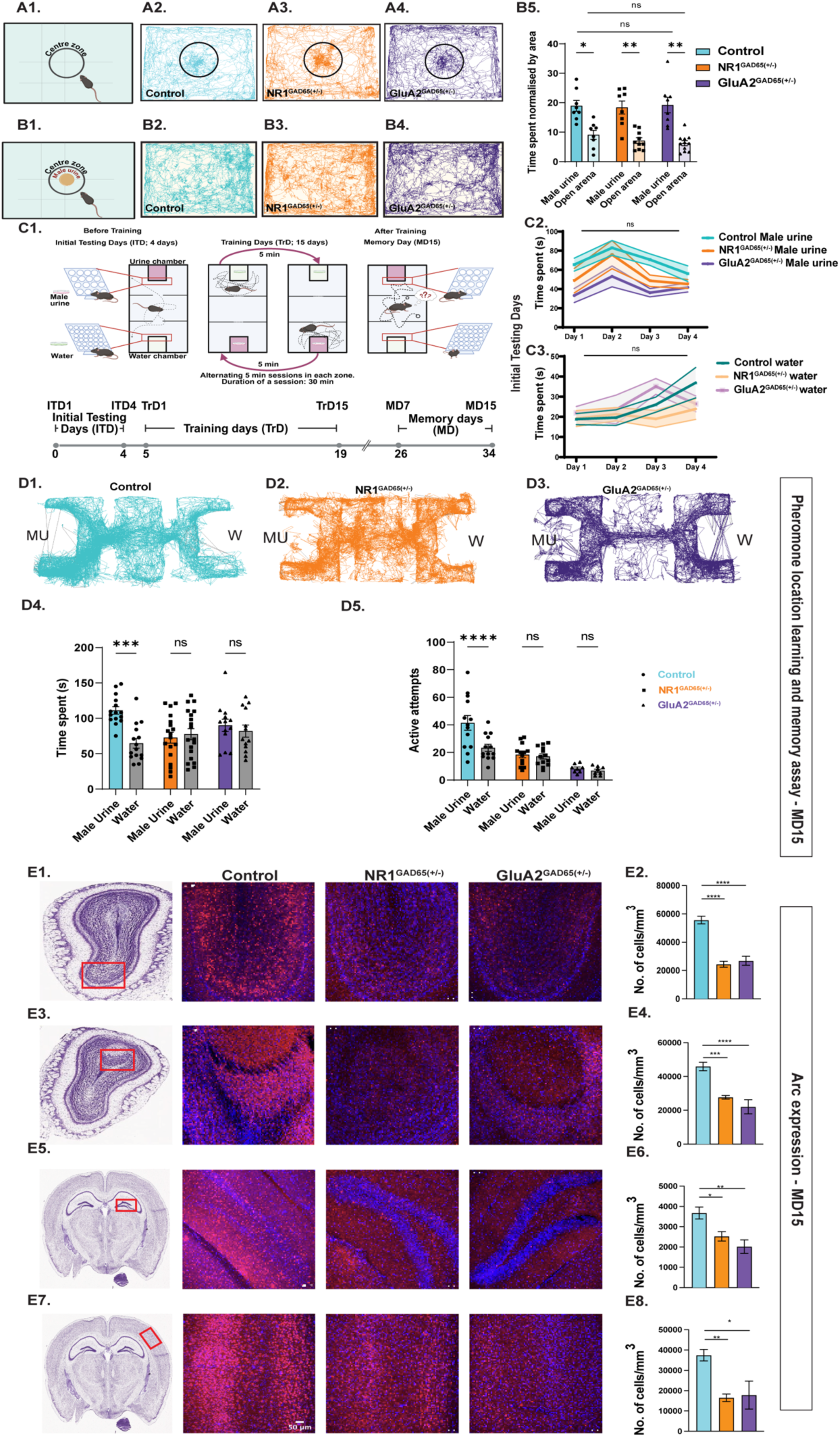
Learning and memorizing pheromone locations require functional NMDARs and AMAPARs in GAD65-expressing interneurons. A1 and B1. Illustrations depicting task design for pheromonal detection and empty chamber center-zone (open arena) exploration. Pheromonal detection readout measured by the time spent by mice to explore pheromone-containing petri dish compared with the empty center-zone in the similar open arena. A2, A3, and A4 Representative path taken by animals while exploring the pheromone-containing petri dish in an open arena. B2, B3, and B4 Representative path taken by animals while exploring the empty center-zone in the similar open arena. B5. Bar-graph comparing time spent in pheromone-containing chamber and open arena (normalized by the area) by different groups of mice. All 3 groups, Control, NR1^GAD65(+/-)^, and GluA2^GAD65(+/-)^ spent significantly higher time exploring the pheromone-containing center zone compared to empty chamber centre-zone (open arena) (Two-way ANOVA, followed by Tukey’s multiple comparison tests, * p < 0.05, ** p < 0.01, *** p < 0.001, **** p < 0.0001, n = 8 controls, n = 8-10 NR1^GAD65(+/-)^ & GluA2^GAD65(+/-)^ 8-11, Control shown in cyan, NR1^GAD65(+/-)^ shown in orange and GluA2^GAD65(+/-)^ shown in purple). C1. Illustration depicting pheromone location learning and memory paradigm and timeline followed for training and memory quantification. C2. Time spent near the male urine chamber by 3 groups of female mice, i.e. Control, NR1^GAD65(+/-)^, and GluA2^GAD65(+/-)^, respectively, during the initial testing days (ITDs) (Two-way ANOVA (mixed-effects model with Geisser–Greenhouse correction), followed by Tukey’s multiple comparisons test showed a significant main effect of Days, *p* = 0.04, and Genotype, p = 0.004. Days × Genotype interaction was not significant, *p* = 0.7. However, in pairwise comparisons, the differences were not observed after the Tukey’s correction for multiple comparison. C3. Time spent near the water chamber by 3 groups of female mice, i.e. Control, NR1^GAD65(+/-)^, and GluA2^GAD65(+/)^, respectively, during the initial testing days (Two-way ANOVA (mixed-effects model with Geisser–Greenhouse correction), showed a non-significant main effect of Days, *p* = 0.18, and Genotype, p = 0.36. Days × Genotype interaction was not significant, *p* = 0.44. D1, D2, and D3. Tracks taken by Control, NR1^GAD65(+/-)^, and GluA2^GAD65(+/)^ female mice on memory day 15 (MD15), obtained by the nose-point feature of EthoVision software (8.5 XT). MU represents the male urine chamber, and W represents the water chamber. D4. Bar graphs showing Time spent near the male urine and water chamber by control, NR1^GAD65(+/-)^, and GluA2^GAD65(+/)^ females on memory day, MD 15 (Two-way ANOVA, followed by Tukey’s multiple comparison test, a significant main effect of stimulus [p (Male Urine vs. Water) = 0.01], was observed, indicating that the time spent investigating urine samples was different from the time spent in water chamber. The effect of genotypes was non-significant, *p* = 0.2. The effect of interaction between Stimulus and Genotype was significant, *p* = 0.03. Pairwise interaction showed Control Male urine vs. Control water, p = 0.0005, p > 0.05 for other pairwise interactions indicated with ns, N_Control_ = 15, N_NR1GAD65(+/-)_ = 19, N_GluA2GAD65(+/-)_ = 13). Controls exhibited significantly higher Time spent near urine chamber compared to water chamber on memory day, MD15, suggesting intact memory of pheromone location. NR1^GAD65(+/-)^ females and GluA2^GAD65(+/-)^ females displayed similar Time spent near both chambers on memory testing day, MD 15, indicating impaired memory. D5. Bar graphs showing Active attempts exhibited by the Control, NR1^GAD65(+/-)^, and GluA2^GAD65(+/-)^ females towards the male urine and water chamber on memory day, MD 15 (Two-way ANOVA, followed by Tukey’s multiple comparison test, there were significant main effects of stimulus [(Male Urine vs. Water) *p* = 0.005] and genotypes (*p* < 0.0001). The interaction between Stimulus × Genotype was also significant, *p* = 0.0003. This show overall differences between Control, NR1^GAD65(+/-),^ and GluA2^GAD65(+/-)^ mice. Pairwise interaction showed Control Male urine vs. Control water p < 0.0001, p > 0.05 for other pairs tested indicated with ns, N_Control_ = 13, N_NR1GAD65(+/-)_ = 13, N_GluA2GAD65(+/-)_ = 8). Controls exhibited significantly higher number of active attempts made towards the urine chamber compared to water chamber on memory day, MD15, suggesting intact memory of pheromone location. NR1^GAD65(+/-)^ and GluA2^GAD65(+/-)^ females showed a similar number of active attempts to both chambers on memory testing day, MD 15, indicating memory impairment. E1. Immunofluorescence images of MOB on MD15 of control, NR1^GAD65(+/-)^ and GluA2^GAD65(+/-)^. (Blue: DAPI, Red: Arc, Scale bar: 50 µm). E2. Significantly lower number of Arc positive cells were observed in the MOB of NR1^GAD65(+/-)^ and GluA2^GAD65(+/-)^ animals compared to control mice (p < 0.0001, F = 29.29, Ordinary one-way ANOVA, Bonferroni’s multiple comparisons test, Control vs NR1^GAD65(+/-)^: p < 0.0001, NR1^GAD65(+/-)^ vs GluA2^GAD65(+/-)^: p > 0.9232, Control vs GluA2^GAD65(+/-)^: p < 0.0001, n = 5). E3. Immunofluorescence images of AOB on MD15 of control, NR1^GAD65(+/-)^ and GluA2^GAD65(+/-)^. E4. Significantly lower number of Arc positive cells were observed in the AOB of NR1^GAD65(+/-)^ and GluA2^GAD65(+/-)^ heterozygous knockout animals compared to control mice (p < 0.0001, F = 16.52, Ordinary one-way ANOVA, Bonferroni’s multiple comparisons test, Control vs NR1^GAD65(+/-)^: p = 0.0007, NR1^GAD65(+/-)^ vs GluA2^GAD65(+/-)^: p = 0.5658, Control vs GluA2^GAD65(+/-)^: p < 0.0001, n = 3). E5. Immunofluorescence images of hippocampus on MD15 of control, NR1^GAD65(+/-)^ and GluA2^GAD65(+/-)^. E6. Significantly lower number of Arc positive cells were observed in the Dentate Gyrus of NR1^GAD65(+/-)^ and GluA2^GAD65(+/-)^ heterozygous knockout animals compared to control mice (p = 0.0016, F = 8.376, Ordinary one-way ANOVA, Bonferroni’s multiple comparisons test, Control vs NR1^GAD65(+/-)^: p = 0.0352, NR1^GAD65(+/-)^ vs GluA2^GAD65(+/-)^: p = 0.6882, Control vs GluA2^GAD65(+/-)^: p = 0.0014, n = 3). E7. Immunofluorescence images of SSC on MD15 of control, NR1^GAD65(+/-)^ and GluA2^GAD65(+/-)^. Significantly lower number of Arc positive cells were observed in the SSC of NR1^GAD65(+/-)^ and GluA2^GAD65(+/-)^ heterozygous knockout animals compared to control mice (p = 0.0027, F = 7.562, Ordinary one-way ANOVA, Bonferroni’s multiple comparisons test, Control vs NR1^GAD65(+/-)^: p = 0.0061, NR1^GAD65(+/-)^ vs GluA2^GAD65(+/-)^: p > 0.9999, Control vs GluA2^GAD65(+/-)^: p = 0.0105, n = 3). * p < 0.05, ** p < 0.01, *** p < 0.001, **** p < 0.0001, ns p > 0.05

Next, we subjected the animals to a pheromone location learning and memory assay. To investigate if the animals show any inherent preference towards any chamber, the initial testing days (ITD) were conducted as explained (see materials and methods, section Multimodal Pheromone Location Learning Paradigm). The preference was quantified based on the time spent in front of the chambers. The post hoc analysis revealed no significant differences in time spent across the genotypes on all testing days (Figure 2C2-3). Following ITD sessions, a 15-day long training day phase (TrD) was conducted. During TrD, we trained the animals to associate the volatiles and non-volatiles with the specific hole-size of the plates. Animals showed consistent attempts to sample the pheromones, quantified by the time spent and active attempts, throughout the TrD, i.e. on training day 1 (TrD1), 7, and 15 (supplementary Figure S2). 15 days after the TrD was over, we quantified the memory of pheromone location, on memory day 15 (MD 15). For testing the memory, we used a new but similar setup to avoid the presence of any pheromonal cues while probing the memory of pheromone locations. We observed an increased preference towards pheromone zone by control females, shown by significantly higher time spent (Figure 2 D4, Two-way ANOVA, followed by Tukey’s multiple comparison test, p = 0.0005, N = 15) and active attempts (Figure 2 D5, Two-way ANOVA, followed by Tukey’s multiple comparison test, p < 0.0001, N = 13). However, GluA2^GAD65(+/-)^ females showed similar time spent (Figure 2 D4, Two-way ANOVA, followed by Tukey’s multiple comparison test, p = 0.98, N = 13) and number of active attempts (Figure 2 D5, Two-way ANOVA, followed by Tukey’s multiple comparison test, ns p = 0.8, N = 8) towards pheromonal and neutral stimulus chambers. Similarly, NR1^GAD65(+/-)^ females showed no significant difference in time spent (Figure 2 D4, Two-way ANOVA, followed by Tukey’s multiple comparison test, p = 0.99, N = 19) and active attempts (Figure 2 D5, Two-way ANOVA, followed by Tukey’s multiple comparison test, p = 0.99, N = 13) made towards any chamber. These results prove the essential role of iGluRs in regulating pheromone location learning and memory. To probe the memory of pheromone chamber at an early time point, we conducted pheromone location learning assay with another set of female mice and checked their memory 7 days post-TrD. We observed a lack of memory on MD7, shown by similar time spent and active attempts towards both chambers (Supplementary Figure S3). Previous studies report the modulation of sniffing during odor discrimination and social communication in animals (31–35). To study if the iGluR modifications cause any differences in the sniffing, we quantified their sniffing toward opposite-sex urine volatiles under head-restrained conditions (8, 29, 32, 36). We did not observe any changes in sniffing frequency across different experimental groups of mice, indicating that the pheromone location learning and memory deficits we report here are not due to the differences in their sampling behaviors (Supplementary Figure S4).

The expression of Arc, an immediate early gene, indicates neural activity during memory recall. Previously, we have seen differential expression of Arc in the OB, somatosensory cortex (SSC), and Dentate Gyrus (DG) of animals displaying a lack of pheromone location memory (8). To understand the neuronal activation pattern during MD15 across groups, we performed immunohistochemistry against Arc antibody. When quantified, we observed a significantly lesser number of Arc+ cells in MOB, AOB, SSC, and DG (Figure 2 E1 – E8: Ordinary one-way ANOVA, Bonferroni’s multiple comparisons test, p < 0.05, n = 5 for control mice and n = 3 for NR1^GAD65(+/-)^ and GluA2 ^GAD65(+/-)^) compared to controls. This decrease in the Arc+ cells in NR1^GAD65(+/-)^ and GluA2 ^GAD65(+/-)^ females on MD15 explains the behavioral phenotypes we observed.

### Ionotropic glutamate receptors in the inhibitory interneurons of accessory olfactory bulb regulate pheromone location learning and memory

Having observed the pheromone location learning and memory deficits in GAD65-expressing interneuron-specific iGluR heterozygous knockouts, we decided to probe the role of inhibitory interneuron network of OB in regulating pheromone location learning and memory. Earlier, we have shown that modulating the inhibitory network can affect olfactory discrimination abilities (15, 18, 37). To modulate the inhibitory network of MOB, we injected AAV5-Cre expressing viral particles in the granule cell layer (GCL) of OB in NR1Lox and GluA2Lox female mice, using a well-established stereotaxic coordinate (15, 37). As we have observed the involvement of AOB neurons in pheromone location learning, we targeted whole of MOB and AOB interneurons in these mice using stereotaxic approaches (Figure 3 A1-A3). In three different groups of mice – control, NR1Lox, and GluA2Lox, the iGluR functions were modulated in the inhibitory network of MOB and AOB. These modified mice are referred as NR1^𝚫OB^, GluA2^𝚫OB^ henceforth. The behavioral training started 2-3 weeks after the surgery. Pheromone detection abilities of the NR1^𝚫OB^ and GluA2^𝚫OB^ animals were comparable to Controls (Figure 3B1-4, 3B4; Ordinary one-way ANOVA, Bonferroni’s multiple comparison tests, p > 0.05). After 15 days of training, we evaluated their memory on MD 15. While we observed enhanced time spent (Figure 3 D1, Two-way ANOVA, followed by Tukey’s multiple comparison test, n = 9, p = 0.02) and number of active attempts (Figure 3 D2, Two-way ANOVA, followed by Tukey’s multiple comparison test, n = 8, p = 0.02) towards the pheromone chamber by the control female mice. NR1^𝚫OB^ group spent similar time (Figure 3 D1, Two-way ANOVA, followed by Tukey’s multiple comparison test, p = 0.99, n = 7) and exhibited similar number of active attempts (Figure 3 D2, Two-way ANOVA, followed by Tukey’s multiple comparison test, p = 0.6, n = 7) towards both the urine and water chamber on MD15. Similarly, GluA2^𝚫OB^ females exhibited no difference in the time spent (Figures 3 D1, Two-way ANOVA, followed by Tukey’s multiple comparison test, p = 0.2 n =8), and active attempts (Figure 3 D2, Two-way ANOVA, followed by Tukey’s multiple comparison test, p = 0.5, n = 8) towards any specific chamber. These results confirm the role of ionotropic glutamate receptors of the OB in regulating the learning and memory of pheromone locations.

**Figure 3:**
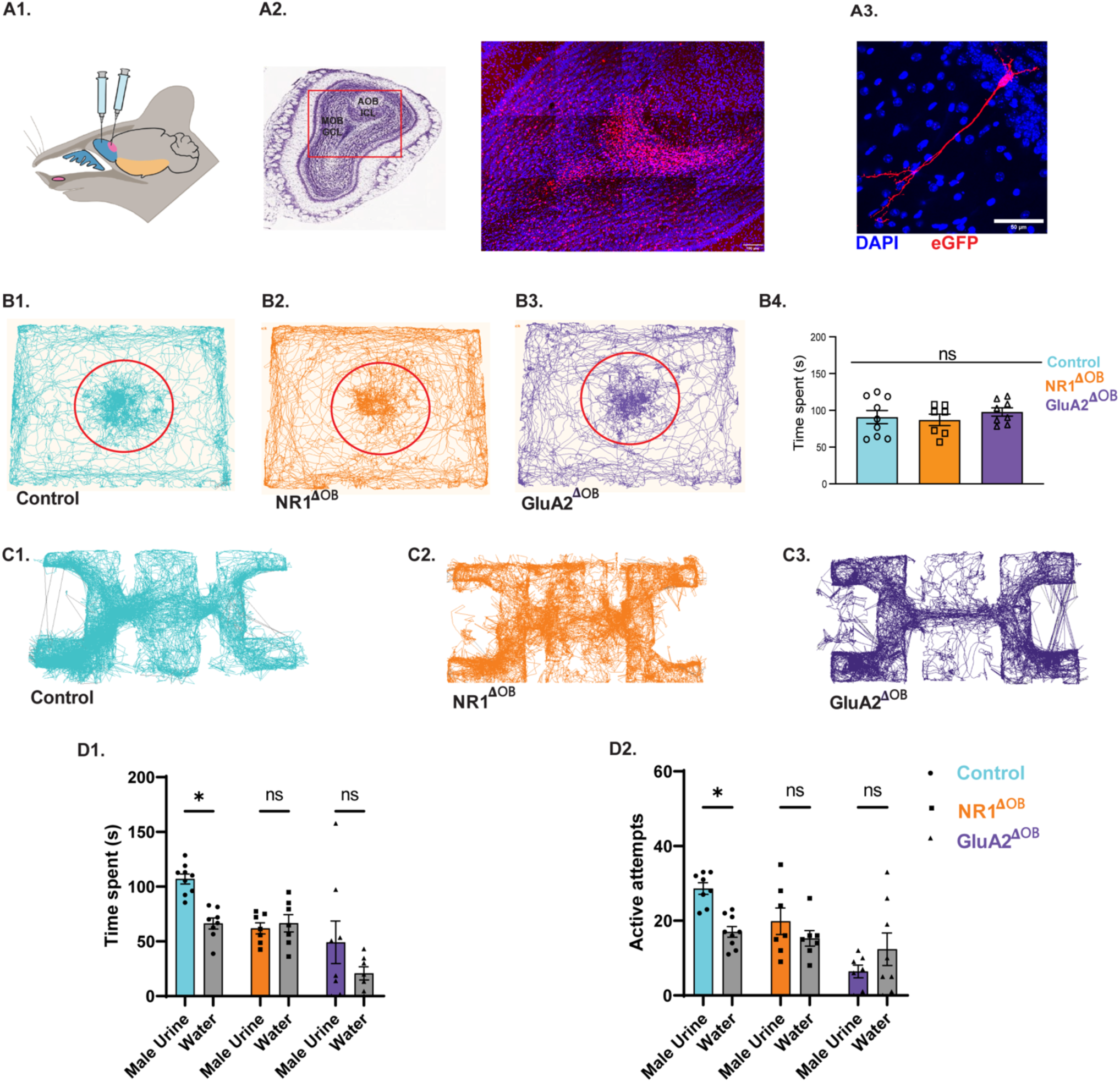
Modification of ionotropic glutamate receptors in the inhibitory interneurons of olfactory bulb results in the deficits of pheromone location learning A1. Illustration representing stereotaxic delivery of AAV particles in the main and accessory olfactory bulb granule cell regions. A2. Representative images showing eGFP expression in the granule cell layer of MOB and AOB three weeks after the stereotaxic injection of cre viral vectors containing GFP. male urine exposure (Blue: DAPI, Red: eGFP, scale bar = 100 µm). A3. MOB GCL interneuron expressing eGFP after AAV-mediated Cre delivery. B1, B2, & B3. Examples of tracks taken by the control, NR1^𝚫OB^, and GluA2^𝚫OB^ animal, analyzed using the “Nose-point” feature of EthoVision software in a pheromone detection assay. As seen by the tracks, different experimental groups spent similar time near petri-dish containing male urine (MU) and soiled bedding (SB) shown in a red circle. B4. Similar time was spent by control, NR1^𝚫OB^, and GluA2^𝚫OB^ female mice in a pheromone detection assay (Ordinary one-way ANOVA, p = 0.6247, F = 0.4811, n_control_ = 9, n_NR1𝚫OB_ = 7, n_GluA2𝚫OB_ = 8). Data are shown in mean ± SEM. C1, C2, & C3. Examples of tracks taken by Control, NR1^𝚫OB^, and GluA2^𝚫OB^ female mice on memory day (MD 15) in pheromone location learning assay. D1. Bar graphs representing Time spent by Control, NR1^𝚫OB^, and GluA2^𝚫OB^ female mice near the male urine and water chamber on the memory day, MD 15, (Two-way ANOVA, followed by Tukey’s multiple comparison test, showed significant main effect of stimulus (Male Urine vs. Water), *p* = 0.009, indicating that the time spent investigating urine samples was different from the time spent in the water chamber. A significant main effect of Genotype was also observed, *p* = 0.002, showing overall differences between Control, NR1^𝚫OB^ and GluA2^𝚫OB^ mice. The interaction between Stimulus × Genotype was also significant *p* = 0.04. Pairwise comparisons showed, Control Male urine vs. Control water p = 0.03, and p > 0.05 for other pairs tested indicated with ns, n_Control_ = 9, n_NR1𝚫OB_ = 7, n_GluA2𝚫OB_ = 8. Control mice spent higher time exploring male urine chamber compared to water chamber on memory day, MD 15, indicating intact memory. NR1^𝚫OB^ and GluA2^𝚫OB^ females exhibited similar Time spent near both urine and water chambers on memory testing day, MD 15 suggesting memory impairment. D2. Bar graph showing number of Active attempts made by Control, NR1^𝚫OB^, and GluA2^𝚫OB^ female mice towards the male urine and water chamber on memory day, MD 15. Two-way ANOVA, followed by Tukey’s multiple comparison test, showed a significant main effect of Genotype, *p* = 0.002, showing overall differences between Control, NR1^𝚫OB^, and GluA2^𝚫OB^ mice. The interaction between Stimulus × Genotype was also significant *p* = 0.002. Pairwise comparisons showed, Control Male urine vs. Control water p = 0.02, and p > 0.05 for other pairs tested indicated with ns, n_Control_ = 9, n_NR1𝚫OB_ = 7, n_GluA2𝚫OB_ = 8). Control mice exhibited higher number of active attempts towards male urine chamber compared to water chamber on memory day, MD 15, indicating intact memory. NR1^𝚫OB^ and GluA2^𝚫OB^ females performed a similar number of active attempts towards both chambers on memory testing day, MD 15, indicating impaired pheromone location learning and memory * p < 0.05, ** p < 0.01, *** p < 0.001, **** p < 0.0001, ns p > 0.05

Mouse AOB plays a critical role in pheromone information processing. To further investigate whether even perturbing the AOB inhibitory network only alters the pheromone location learning abilities, we kept the MOB iGluRs intact, and modified the AOB circuits. We injected AAV5-Cre viral particles in the AOB of NR1Lox and GluA2Lox female animals. They are referred to as NR1^𝚫AOB^ and GluA2^𝚫AOB^ henceforth. After three weeks of surgery, their pheromone detection abilities were assessed and compared with that of control mice. NR1^𝚫AOB^ and GluA2^𝚫AOB^ female mice showed undisturbed pheromone detection abilities, comparable to controls as observed by the time spent near a male urine-containing petri dish (Figure 4B1-B4; B4: Ordinary one-way ANOVA, Bonferroni’s multiple comparison tests, p > 0.05, n = 7-9). Mice were then trained to associate specific hole sizes with a pheromone-containing chamber and neutral stimulus chamber for 15 days. After 15 days, their memory was evaluated. Control mice showed enhanced time spent (Figure 4 D1, Two-way ANOVA, followed by Tukey’s multiple comparisons test, p < 0.0001, n = 9) and number of active attempts towards the pheromone chamber compared to the neutral stimulus chamber (Figure 4 D2, Two-way ANOVA, followed by Tukey’s multiple comparisons test, p < 0.002, n = 9). However, similar time spent and number of active attempts were made by NR1^𝚫AOB^, GluA2^𝚫AOB^ females compared to control animals (Figure 4 D1 and D2, Two-way ANOVA, followed by Tukey’s multiple comparisons test, D1; NR1^𝚫AOB^,p = 0.99, n = 7, GluA2^𝚫AOB^, p = 0.6, n = 8, D2; NR1^𝚫AOB^, p = 0.73, n = 7, GluA2^𝚫AOB^, p = 0.96, n = 8). These results confirm the role of ionotropic glutamate receptors of the AOB in regulating the learning and memory of pheromone locations. Our results thus prove the important role of iGluRs on GAD65 expressing interneurons both in MOB and AOB, and thereby the synaptic inhibitory network in regulating the pheromone location learning and memory.

**Figure 4:**
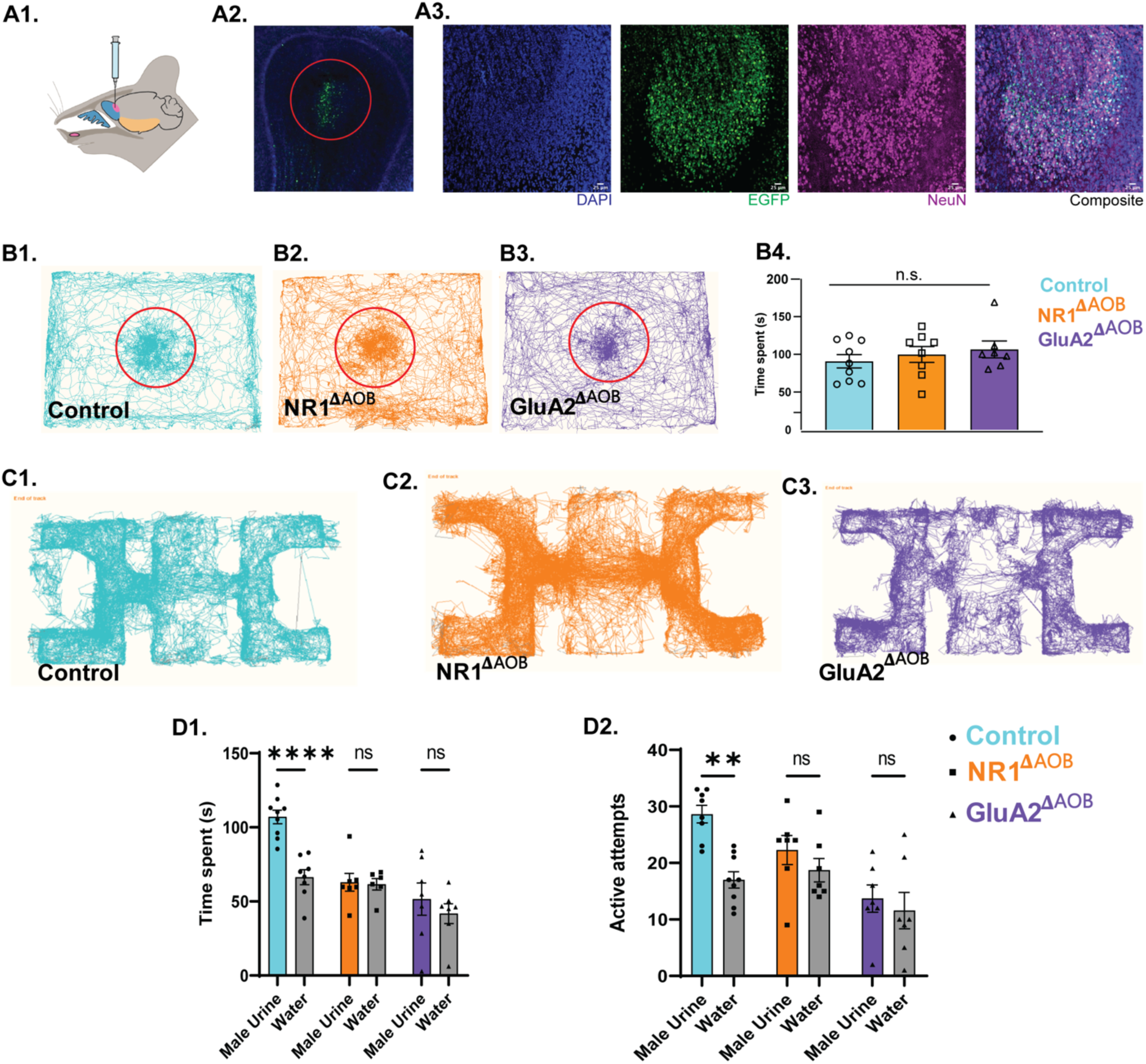
Ionotropic glutamate receptor functions in the accessory olfactory bulb neuronal circuitry regulate pheromone location learning A1. Diagrammatic representation of stereotaxic delivery of AAV particles in the accessory olfactory bulb. A2. Expression of EGFP as an indication of AAV-mediated Cre expression in AOB. A3. Confocal images of Cre-viral particles (Green) injected section co-stained with NeuN (Magenta) and DAPI (Blue) (Scale bar: 25 μm). B1, B2, and B3. Examples of tracks taken by the Control, NR1^𝚫AOB^, and GluA2^𝚫AOB^ female mice, analyzed using the “Nose-point” feature of EthoVision software in a pheromone detection assay. As seen by the tracks, different experimental groups spent similar time near petri-dish containing male urine and soiled bedding shown in a red circle. B4. Similar time was spent by the Control, NR1^𝚫AOB^, and GluA2^𝚫AOB^ female mice in a pheromone detection assay (Ordinary one-way ANOVA, p = 0.6139, F = 0.5507, n_Control_ = 9, n_NR1𝚫AOB_ = 8, n_GluA2𝚫AOB_ = 7). Data are mean ± SEM. (). C1, C2, and C3. Examples of tracks taken by the Control, NR1^𝚫AOB^, and GluA2^𝚫AOB^ female mice on memory day, MD 15, in pheromone location learning assay. D1. Bar graphs showing Time spent by the Control, NR1^𝚫AOB^, and GluA2^𝚫AOB^ female mice near the male urine and water chamber on memory day, MD 15 (Two-way ANOVA, followed by Tukey’s multiple comparisons test showed a significant main effect of stimulus (Male Urine vs Water), *p* = 0.002, indicating that the time spent investigating urine samples was different from the time spent in the water chamber. A significant main effect of Genotype was also observed, *p* = 0.0001, showing overall differences among Control, NR1^𝚫AOB^ and GluA2^𝚫AOB^ female mice. The interaction between Stimulus × Genotype was also significant *p* = 0.008. Pairwise comparison showed Control Male urine vs Control water p < 0.0001, and p > 0.05 for other pairs tested indicated with ns, n_Control_ = 9, n_NR1𝚫AOB_ = 7, n_GluA2𝚫AOB_ = 7). Control mice spent significantly higher time near male urine chamber compared to water chamber on memory day, MD 15, indicating intact memory. However, NR1^𝚫AOB^ and GluA2^𝚫AOB^ females exhibited memory impairment, as they spent a similar amount of time near both chambers on memory day, MD 15. D2. Bar graphs showing Active attempts exhibited by the Control, NR1^𝚫AOB^, and GluA2^𝚫AOB^female mice towards the male urine and water chamber on memory day, MD 15 (Two-way ANOVA, followed by Tukey’s multiple comparisons test showed a significant main effect of stimulus, *p* = 0.004, showing overall differences between Control, NR1^𝚫AOB^, and GluA2^𝚫AOB^ mice. A significant main effect of Genotype was also observed, *p* = 0.003. Pairwise comparisons showed Control Male urine vs Control water p < 0.002, and p > 0.05 for other pairs tested indicated as ns, n_Control_ = 9, n_NR1𝚫AOB_ = 7, n_GluA2𝚫AOB_ = 7. Control mice exhibited higher number of active attempts towards male urine chamber compared to water chamber on memory day, MD 15, indicating intact memory. However, NR1^𝚫AOB^ and GluA2^𝚫AOB^ females performed a similar number of active attempts towards both chambers on memory testing day, MD 15, indicating impaired pheromone location learning and memory. * p < 0.05, ** p < 0.01, *** p < 0.001, **** p < 0.0001, ns p > 0.05

## Discussion

Rodents’ olfactory system plays a major role in socio-sexual communication (38, 39). Urinary scent marks, containing volatile and non-volatile pheromones, are essential for such communication (9, 40). The volatile components of urine can reflect the reproductive potential of a male mouse as their production is testosterone-dependent (41). Therefore, the reproductive success of a female mouse would be controlled by their ability to remember the pheromone locations in the wild. As the pheromones are streaked on objects of different sizes and shapes, the involvement of the somatosensory system is expected in this process. We made use of these observations to design our behavioral paradigm for studying the neural mechanisms of pheromone location learning and memory (8, 42). Further, our study took advantage of the technique to spatio-temporal deletion of iGluRs from the inhibitory network of MOB and AOB by using the Cre-Lox system and viral vector-mediated gene delivery (15).

Both main and accessory olfactory bulbs are involved in pheromonal information processing (43, 44). Previous work from the lab validates the behavioral assay we used in the present study as an ideal paradigm for investigating the engagement of olfactory and whisker systems in pheromone location learning and memory (8). Therefore, we decided to modify the MOB and AOB circuits and probe animals’ ability to learn and memorize pheromone locations using this paradigm. On training mice in this pheromonal location learning assay, we observed many c-Fos positive cells, a neuronal activity marker, among the interneurons of both MOB and AOB circuits. Increased c-fos expression in the AOB after the pheromone location learning assay, indicates an increase in activation of inhibitory interneurons after repeated sensory experience. Different studies utilizing activity-dependent marker expression pattern analysis have demonstrated an increase in the maturation and recruitment of adult-born interneurons in the AOB following social encounter or exposure to relevant social cues (45, 46). Further, to confirm the activation of inhibitory network, we expressed GCaMP6f driven by GAD-65, which is expressed by the majority of the inhibitory interneurons of MOB and AOB (17, 47). AOB interneurons showed higher calcium activity in response to urinary volatiles compared to airflow stimuli in both untrained and trained conditions (Figure 1). This provides direct, population-level evidence that AOB interneurons are dynamically engaged during the association task. These findings reflect activity-dependent recruitment of inhibitory populations in the AOB circuitry during associative learning, which likely enhances the signal-to-noise ratio and refines pheromonal processing. While cell-type-specific contributions remain an important direction for further work, these findings provide a necessary foundation for understanding how overall AOB inhibition regulates pheromone location learning and plasticity.

To modulate the inhibitory actions, we targeted iGluRs of GAD65-expressing interneurons, and their deletion did not result in any anxiety-and depressive-like symptoms (Figure S1). While the heterozygous knockouts of GluA2 (subunit of AMPARs) and NR1 (subunit of NMDARs) showed intact pheromone detection abilities, their pheromone location memory was reduced compared to control mice. There was a correlated reduction in the number of Arc-expressing cells in the knockout mice (Figure 2). Further, to make the perturbation specific to olfactory areas, interneurons of both MOB & AOB, and AOB alone were targeted by making use of Cre-Lox system and stereotaxic delivery of viral vectors. In both these cases pheromone location memory was compromised compared to control animals. This reveals the central role of iGluRs in MOB and AOB inhibitory networks in regulating pheromone location learning and memory (Figures 3 and 4). Our use of the region-specific stereotaxic approach ensured that the observed behavioral and physiological changes are consistent with the modulation of local inhibitory signaling in the AOB and MOB rather than off-target effects.

To study the neuronal mechanisms, it is critical to form ethologically relevant associations in a learning paradigm. The urinary scent marks communicate the competitive ability and identity of the male mouse (48, 49). Therefore, to mimic the natural conditions, we designed the experimental paradigm, where the animals had to use both olfactory and whisker systems during learning. The whisker system can potentially be involved while sniffing the pheromones streaked on different objects. Earlier studies report coordinated action of sniffing and whisking, which is regulated by respiratory centers in the ventral medulla (50–52). Thus, the female mice might be using this multimodal sampling strategy while exploring pheromone locations. Earlier experiments by Garcia proved the relevance of association when two stimuli are combined. The association of tasty water with a noxious drug worked better compared to a bright-noisy water-noxious stimulus association in avoiding a gustatory stimulus (53). Our paradigm, where animals had to learn to associate certain hole sizes with pheromonal cues, worked well, and mice exhibited a robust memory of this association (Figure 2).

Odorants evoke specific spatiotemporal glomerular activity patterns in the olfactory bulb, which is the first relay station in olfactory information processing (31, 32, 54–56). The overlapping patterns of activity evoked by complex odorants are refined and decorrelated by the inhibitory network of OB before sending the odor information to higher centers (15, 37). We observed significantly higher activated cells among the inhibitory network of MOB and AOB in response to urinary volatiles (Figure 1). Therefore, we modulated the synaptic inhibition by targeting iGluRs on GAD65-expressing interneurons (Figure S1). While removing the GluA2 subunit of AMPARs increases the inflow of calcium, deleting the essential subunit of NMDARs would result in non-functional NMDARs, which may cause a decrease in Ca^2+^ levels. These modifications trigger a bidirectional shift in the inhibition in the OB network (15). Alteration in the optimal inhibition of the OB circuits, results in an impairment in odor discrimination abilities of mice (29, 37). Vomeronasal and main olfactory systems are more integrated in processing information from volatile and non-volatile urinary pheromones (11, 57–59). Therefore, to study the neural mechanisms, we undertook a genetic approach to modify iGluRs present on the GAD65-expressing interneurons in both MOB and AOB and probed the pheromone location memory of these iGluR-modified mice after training. They did not show the memory of pheromone locations compared to control animals (Figure 2). The learning and memory deficit was independent of their sampling behaviors (Supplementary Figure S4).

Deleting GluA2 or NR1 from inhibitory interneurons would result in a global modulation of inhibitory actions, considering the extensive lateral connectivity in MOB and AOB (24, 60–63). Apart from this, there are other sources of glutamatergic inputs on inhibitory neurons, for example, on granule cells (64). As synaptic inhibition has been shown to be critical for decorrelating the overlapping patterns of activity in MOB and the sex-specific cue selectivity of AOB neurons, we decided to make our modulations specific to MOB and AOB circuits by using stereotaxic approaches (24, 37). We used a well-established stereotaxic coordinate to target the MOB interneurons; for the AOB, we optimized the coordinates (15, 30, 37). Modulating iGluRs in MOB and AOB inhibitory circuits resulted in pheromone location learning and memory deficits (Figure 3 and 4). This can be explained by the reports on the role of iGluRs in long-term potentiation and depression, the mechanisms that underly learning and memory (65, 66).

Pheromones can promote associative learning (67). The non-volatile urinary proteins from males attract naïve females. These proteins not only attract females but also promote the learning of urinary volatiles (68, 69). As we kept the bedding with urinary proteins from males in front of the pheromone chamber, from which the urinary volatiles emanated, it facilitated associative learning, as reflected in the memory readouts of all control mice used in the study. This was hampered in case of iGluR modification in the AOB inhibitory circuits (Figures 2 & 4). These results confirm the importance of AOB inhibitory circuits in regulating pheromone location learning and memory. The synaptic activation of neurons leads to the expression of immediate early genes, for example, Activity-regulated cytoskeleton-associated (Arc) protein. Newly formed Arc mRNA is accumulated at the strongly activated synapses. The localization of Arc mRNA to active synapses and their transcription requires intact functioning of NMDARs and AMPARs (70, 71). Therefore, iGluR modulations will lead to compromised Arc protein expression, which was observed in iGluR heterozygous knockouts (Figure 2).

Karlson and Luscher defined pheromones as “airborne chemical signals released by an individual into the environment and affecting the physiology and behavior of other members of the same species” (72). Although humans lack neuronal elements in VNO, activation of hypothalamus has been observed in response to pheromone-like compounds (73–75). Further evidence is present for the perception and behavioral changes caused by pheromones in human subjects. For example, synchronization of menstrual cycles and its effect on ovulation in roommates, probably mediated by sweat (76), sniffing sex-steroid-derived compounds resulting in mood changes and arousal (77), social odorants causing modulations in mood (78), etc. All these observations confirm chemical communication in humans, leaving the involvement of olfactory system and underlying neural mechanisms as open questions. As olfactory functions are affected under many disease conditions (79–84), and in long COVID (85–88), investigating the neural mechanisms of chemical communication using appropriate animal models and experimental methods becomes indispensable (8, 89–91).

## Acknowledgments

We thank Laboratory of Neural Circuits and Behavior (LNCB) members and IISER-Pune Biology colleagues for fruitful discussions. We thank staff of *National Facility for Gene Function in Health and Disease* (NFGFHD) and IISER Biology-Leica microscopy facility for the technical support. Some of the illustrations were created with BioRender.com.

## Funding

This work was supported by the DBT/Wellcome Trust India Alliance senior grant (IA/S/22/2/506517 to N.M.A.), DBT/Wellcome Trust India Alliance intermediate grant (IA/I/14/1/501306 to N.M.A.), DST-Cognitive Science Research Initiative (DST/CSRI/2017/271 to N.M.A.), CSIR Fellowship (S.D.M., S.P., S.D.), and UGC Fellowship (D.R.). Part of the work was carried at the National Facility for Gene Function in Health and Disease (NFGFHD) at IISER Pune, supported by a grant from the Department of Biotechnology, Govt. of India (BT/INF/22/SP17358/2016).

## Author Contributions

N.M.A. supervised all aspects of the project. N.M.A. carried out the study conceptualization and experimental design. S.D.M. performed behavioral assays, stereotaxic modulations, and immunohistochemistry experiments and analyzed the data. S.P., D.R., and S.D. performed calcium imaging experiments and analyzed the data.

N.M.A. and S.D.M. wrote the manuscript.

## Competing Interests

The authors have stated explicitly that there are no conflicts of interest in connection with this article.

## Data and Materials Availability

All cumulative data are available in the article/supplementary materials; further inquiries can be directed to the corresponding author.

## List of supplementary material

Supplementary Figures S1 to S4

## Notes

### Competing Interest Statement

The authors have declared no competing interest.

### Summary of Updates

To validate the involvement of GAD65-expressing interneurons in processing pheromonal cues and pheromonal location learning, we conducted calcium imaging experiments using transgenic female mice expressing GCaMP6f in GAD65-expressing interneurons, which comprise majority of the interneuron population in MOB and AOB, before and after training (Figure 1B). The GRIN lens was implanted in the AOB of GAD65-GCaMP6f transgenic female mice (Figure 1 B4). As a control, calcium activity was also recorded in response to airflow used to deliver pheromonal volatiles (0.4 liter per minute, LPM). This population activity was measured in awake, head-restrained condition. The fluorescence change (Delta F/F0) was quantified for 40 trials across 5 females in untrained and trained conditions (Figure 1 B3-B6). Calcium responses were observed from the interneuron population, indicating their involvement in processing the pheromonal information (Figure 1 B5-B6). Further, on quantifying the calcium responses post-training day 15, we observed a marked increase selectively for the male urine but not for airflow stimulation, proving the involvement of AOB inhibitory network in pheromone location learning and memory (Figure 1 B7).

