## Supplemental Figures for "Synaptic inhibition in the accessory olfactory bulb regulates pheromone location learning and memory"

**Supplementary Material:**

**Supplementary Figures:**

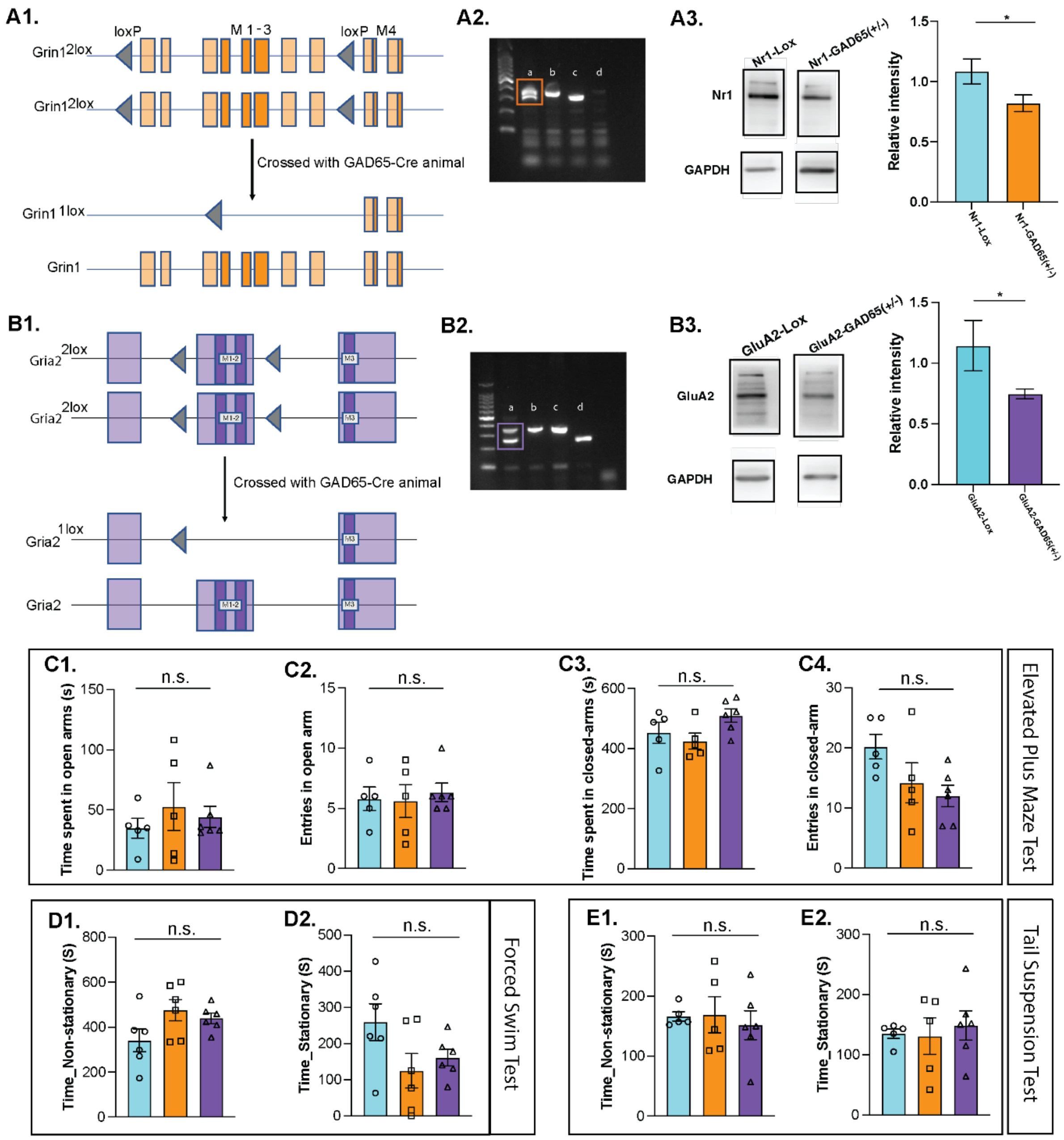


**Supplementary Figure S1: NMDA and AMPA receptor perturbations did not cause any despair-like and anxiety-like behaviors**

A1. Scheme of genetic cross used to generate NR1$^{GAD65(+/-)}$ animals.

A2. Agarose gel electrophoresis image of DNA samples taken from a: Heterozygous NR1$^{GAD65(+/-)}$ animal (369 bp and 315 bp), b: NR1-2Lox (369 bp), c: wild-type (315 bp) and d: Blank.

A3. Immunoblots showing expression of NR1 in the olfactory bulb of control and NR1$^{GAD65(+/-)}$ mice. GAPDH was used as a loading control. NR1 expression was reduced in NR1$^{GAD65(+/-)}$ animals. (Control: 1.083 ± 0.1049, NR1$^{GAD65(+/-)}$: 0.8197 ± 0.0688 mean ± SEM, Unpaired one-tailed t-test, p = 0.0313, n = 6).

B1. Scheme of genetic cross used to generate GluA2$^{GAD65(+/-)}$ animals.

B2. Agarose gel electrophoresis image of DNA samples taken from a: Heterozygous GluA2$^{GAD65(+/-)}$ KO animal (348 bp and 254 bp), b: GluA2-2Lox (348 bp), c: wild-type (254 bp) and d: Blank.

B3. Immunoblots showing expression of GluA2 in the olfactory bulb of control and GluA2$^{GAD65(+/-)}$ mice. GAPDH was used as a loading control. GluA2 expression was reduced in GluA2$^{GAD65(+/-)}$ animals (control: 1.027 ± 0.2051, GluA2: 0.745 ± 0.0400, mean ± SEM Unpaired one-tailed t-test, $p = 0.0347$, $n = 6$).

Elevated Plus Maze Test (EPMT):

C1. Time spent in the open arms by different groups of mice were similar ($p = 0.6437$, $F = 0.4558$, ordinary one-way ANOVA, Bonferroni's multiple comparison tests, control vs. $NR1^{GAD65(+/-)}$, control vs. $GluA2^{GAD65(+/-)}$, and $NR1^{GAD65(+/-)}$ vs. $GluA2^{GAD65(+/-)}$: $p > 0.9999$, control and $NR1^{GAD65(+/-)}$ $n = 5$, $GluA2^{GAD65(+/-)}$ $n = 6$) (Control shown in cyan, $NR1^{GAD65(+/-)}$ shown in orange and $GluA2^{GAD65(+/-)}$ shown in purple).

C2. Time spent in the closed arms by different groups of mice were similar ($p = 0.1158$, $F = 2.556$, ordinary one-way ANOVA, Bonferroni's multiple comparison tests, control vs. $NR1^{GAD65(+/-)}$: $p > 0.9999$, control vs. $GluA2^{GAD65(+/-)}$ $p = 0.4935$, $NR1^{GAD65(+/-)}$ vs. $GluA2^{GAD65(+/-)}$: $p = 0.1402$).

C3. Number of entries in open arms across groups were similar ($p = 0.8688$, $F = 0.1422$, ordinary one-way ANOVA, Bonferroni's multiple comparison tests, control vs. $NR1^{GAD65(+/-)}$, control vs. $GluA2^{GAD65(+/-)}$ and $NR1^{GAD65(+/-)}$ vs. $GluA2^{GAD65(+/-)}$: $p > 0.9999$).

C4. Number of entries in close arms across groups were similar ($p = 0.0786$, $F = 3.113$, ordinary one-way ANOVA, Bonferroni's multiple comparison tests, control vs. $NR1^{GAD65(+/-)}$: $p = 0.3324$, control vs. $GluA2^{GAD65(+/-)}$ $p = 0.0889$, $NR1^{GAD65(+/-)}$ vs. $GluA2^{GAD65(+/-)}$: $p > 0.9999$).

Forced Swim Test (FST):

D1. There were no significant differences in the time spent mobile across groups ($p = 0.1006$, $F = 2.687$, ordinary one-way ANOVA, Bonferroni's multiple comparison tests, control vs. $NR1^{GAD65(+/-)}$: $p = 0.1219$, control vs. $GluA2^{GAD65(+/-)}$ $p = 0.3684$, $NR1^{GAD65(+/-)}$ vs. $GluA2^{GAD65(+/-)}$: $p > 0.9999$, control and $NR1^{GAD65(+/-)}$ $n = 5$, $GluA2^{GAD65(+/-)}$ $n = 6$).

D2. There were no significant differences in the time spent immobile across groups ($p = 0.1006$, $F = 2.687$, ordinary one-way ANOVA, Bonferroni's multiple comparison tests, control vs. $NR1^{GAD65(+/-)}$: $p = 0.1219$, control vs. $GluA2^{GAD65(+/-)}$ $p = 0.3684$, $NR1^{GAD65(+/-)}$ vs. $GluA2^{GAD65(+/-)}$: $p > 0.9999$).

Tail Suspension Test (TST):

E1. There were no significant differences in the time spent mobile across groups ($p = 0.8402$, $F = 0.1765$, ordinary one-way ANOVA, Bonferroni's multiple comparison tests, control vs. $NR1^{GAD65(+/-)}$, control vs. $GluA2^{GAD65(+/-)}$, and $NR1^{GAD65(+/-)}$ vs. $GluA2^{GAD65(+/-)}$: $p > 0.9999$, control and $NR1^{GAD65(+/-)}$ $n = 5$, $GluA2^{GAD65(+/-)}$ $n = 6$).

E2. There were no significant differences in the time spent immobile across groups ($p = 0.8474$, $F = 0.1677$, ordinary one-way ANOVA, Bonferroni's multiple comparison tests, control vs. $NR1^{GAD65(+/-)}$, control vs. $GluA2^{GAD65(+/-)}$, and $NR1^{GAD65(+/-)}$ vs. $GluA2^{GAD65(+/-)}$: $p > 0.9999$).

Controls are shown in cyan, $NR1^{GAD65(+/-)}$ animals are shown in orange, and $GluA2^{GAD65(+/-)}$ animals are shown in purple. Data are shown in mean ± SEM.

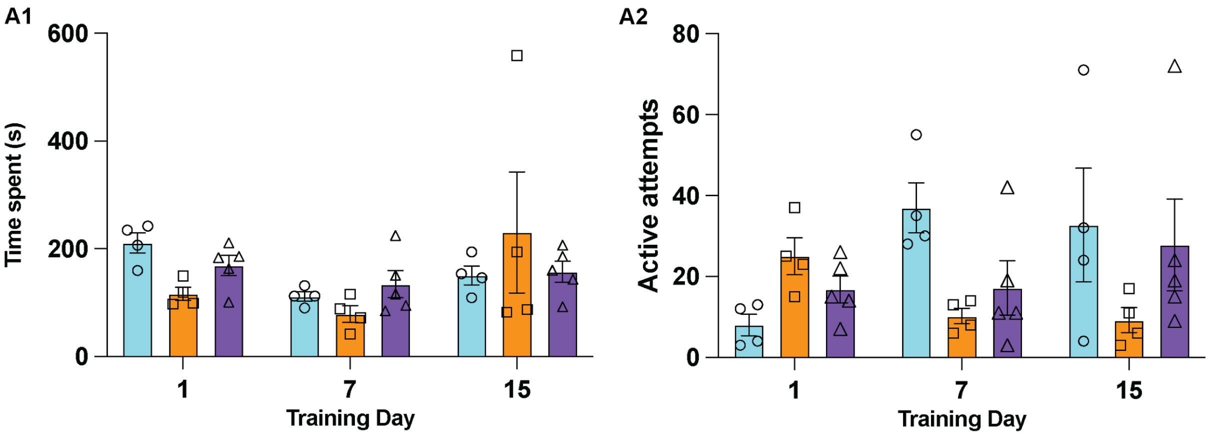


**Supplementary Figure S2: Female mice across different experimental groups showed similar behavior during training sessions.**

A1. Animals across experimental groups displayed similar time spent on TrD 1, 7 and 15 (Two-way ANOVA, Bonferroni's multiple comparison test, p-value > 0.05 across all the groups).

A2. Number of active attempts done by the animals across groups were comparable on TrD 1, 7 and 15 (Two-way ANOVA, Bonferroni's multiple comparison test, p-value > 0.05 across all the groups).

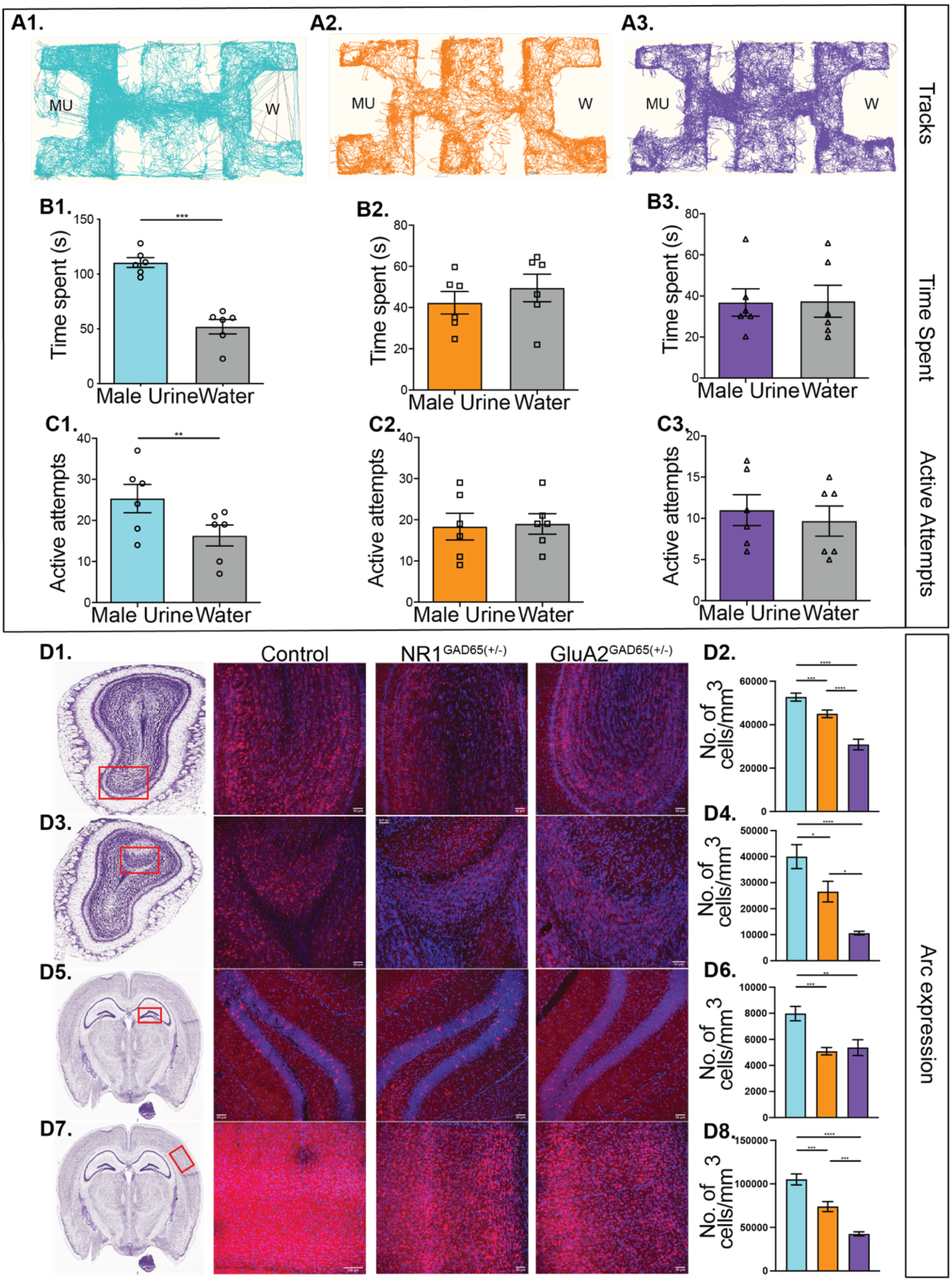


**Supplementary Figure S3: Lack of pheromone location memory in iGluR knockout animals on post-training day 7 (MD7).**

A1, A2, A3: Tracks obtained by the nose-point feature of EthoVision software (8.5 XT) during the memory testing day. MU represents the male urine chamber, and W represents the water chamber.

B1. Significantly longer time was spent near the male urine chamber by control females on memory testing day, MD 7 ($p = 0.0003$, paired two-tailed student's t-test, N = 6 females).

B2. NR1$^{GAD65(+/-)}$ females showed similar time spent near both the chambers on memory testing day, MD 7 (p = 0.3655, paired two-tailed student's t-test, N = 6 females).
B3. GluA2$^{GAD65(+/-)}$ females displayed similar time spent near both the chambers on memory testing day, MD 7 (p = 0.9310, paired two-tailed student's t-test, N = 6 females).
C1. Control female mice exhibited a greater number of active attempts to the male urine chamber on memory testing day, MD 7 (p = 0.0028, paired two-tailed student's t-test, N = 6 females).
C2. NR1$^{GAD65(+/-)}$ females displayed similar number of active attempts to both chambers on memory testing day, MD 7 (p = 0.7866, paired two-tailed student's t-test, N = 6 females).
C3. GluA2$^{GAD65(+/-)}$ females displayed a similar number of active attempts to both chambers on memory testing day, MD 7 (p = 0.2213, paired two-tailed student's t-test, N = 6 females).
D1. Immunofluorescence images of MOB on MD07 of control, NR1$^{GAD65(+/-)}$ and GluA2$^{GAD65(+/-)}$. (Blue: DAPI, Red: Arc, Scale bar: 50 µm).
D2. Significantly lower number of Arc positive cells in MOB of NR1$^{GAD65(+/-)}$ and GluA2$^{GAD65(+/-)}$ heterozygous knockout animals (p < 0.0001, F = 29.77, Ordinary one-way ANOVA, Bonferroni's multiple comparisons test, Control vs NR1$^{GAD65(+/-)}$ : p = 0.0283, NR1$^{GAD65(+/-)}$ vs GluA2$^{GAD65(+/-)}$ : p < 0.0001, Control vs GluA2$^{GAD65(+/-)}$ : p < 0.0001, N = 3).
D3. Immunofluorescence images of AOB on MD07 of control, NR1$^{GAD65(+/-)}$ and GluA2$^{GAD65(+/-)}$ females.
D4. Significantly lower number of Arc positive cells in AOB of NR1$^{GAD65(+/-)}$ and GluA2$^{GAD65(+/-)}$ heterozygous knockout animals (p < 0.0001, F = 14.21, Ordinary one-way ANOVA, Bonferroni's multiple comparisons test, Control vs NR1$^{GAD65(+/-)}$ : p = 0.0476, NR1$^{GAD65(+/-)}$ vs GluA2$^{GAD65(+/-)}$ : p =0.0235, Control vs GluA2$^{GAD65(+/-)}$ : p < 0.0001, N = 3).
D5. Immunofluorescence images of hippocampus on MD07 of control, NR1$^{GAD65(+/-)}$ and GluA2$^{GAD65(+/-)}$.
D6. Significantly lower number of Arc positive cells in the hippocampus of NR1$^{GAD65(+/-)}$ and GluA2$^{GAD65(+/-)}$ heterozygous knockout animals (p = 0.0002, F = 11.55, Ordinary one-way ANOVA, Bonferroni's multiple comparisons test, Control vs NR1$^{GAD65(+/-)}$: p = 0.0004, NR1$^{GAD65(+/-)}$ vs GluA2$^{GAD65(+/-)}$ : p > 0.9999, Control vs GluA2$^{GAD65(+/-)}$ : p = 0.0032, N = 3).
D7. Immunofluorescence images of SSC on MD07 of control, NR1$^{GAD65(+/-)}$ and GluA2$^{GAD65(+/-)}$.
D8. Significantly lower number of Arc positive cells in SSC of NR1$^{GAD65(+/-)}$ and GluA2$^{GAD65(+/-)}$ heterozygous knockout animals (p < 0.0001, F = 35.39, Ordinary one-way ANOVA, Bonferroni's multiple comparisons test, Control vs NR1$^{GAD65(+/-)}$ : p = 0.0006, NR1$^{GAD65(+/-)}$ vs GluA2$^{GAD65(+/-)}$: p = 0.0002, Control vs GluA2$^{GAD65(+/-)}$ : p < 0.0001, N = 3).

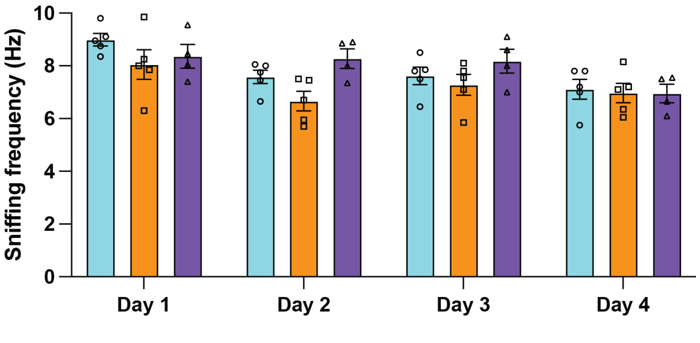


**Supplementary Figure S4: Sniffing frequency towards male urine is unaltered in knockout animals**
Sniffing frequency was similar across four days in all the groups. (p = 0.3552, F = 1.164, Ordinary one-way ANOVA, Bonferroni's multiple comparison tests, control vs. NR1$^{GAD65(+/-)}$: p = 0.7955, NR1$^{GAD65(+/-)}$ vs. GluA2$^{GAD65(+/-)}$: p = 0.5654, control vs. GluA2$^{GAD65(+/-)}$ p > 0.9999, control, NR1$^{GAD65(+/-)}$ N = 5, GluA2$^{GAD65(+/-)}$ N = 4). Controls are shown in cyan, NR1$^{GAD65(+/-)}$ animals are shown in orange, and GluA2$^{GAD65(+/-)}$ animals are shown in purple. Data are shown in mean ± SEM.
